# The *Streptococcus agalactiae* LytSR two-component regulatory system promotes vaginal colonisation and virulence *in vivo*

**DOI:** 10.1101/2024.08.02.606384

**Authors:** Hajar AlQadeeb, Murielle Baltazar, Adrian Cazares, Tiraput Poonpanichakul, Morten Kjos, Neil French, Aras Kadioglu, Marie O’Brien

**Affiliations:** Department of Medical Laboratory Sciences, College of Applied Medical Sciences, Prince Sattam Bin Abdulaziz University, Al-Kharj 11942, Saudi Arabia; Department of Clinical Infection, Microbiology and Immunology, University of Liverpool, UK; Wellcome Sanger Institute, Wellcome Genome Campus, Hinxton, UK; Chakri Naruebodindra Medical Institute, Faculty of Medicine Ramathibodi Hospital, Mahidol University, Thailand; Faculty of Chemistry, Biotechnology and Food Science, Norwegian University of Life Sciences, Ås, Norway; ReNewVax Ltd, UK

**Keywords:** S*treptococcus agalactiae*, Group B *Streptococcus*, newborn infections, LytSR two-component regulatory system, CC-17, pathogenesis

## Abstract

*Streptococcus agalactiae* (or Group B *Streptococcus*, GBS) is a leading cause of neonatal sepsis and meningitis globally. To sense and respond to variations in its environment, GBS possesses multiple two-component regulatory systems (TCSs) such as LytSR. Here, we aimed to investigate the role of LytSR in GBS pathogenicity. We generated an isogenic *lytS* knockout mutant in a clinical GBS isolate and used a combination of phenotypic in vitro assays and in vivo murine models to investigate the contribution of *lytS* to the colonisation and invasive properties of GBS. Deletion of the *lytS* gene in the GBS chromosome resulted in significantly higher survival rates in mice during sepsis, accompanied by reduced bacterial loads in blood, lung, spleen, kidney and brain tissue compared to infection with the wild-type strain. In a mouse model of GBS vaginal colonisation, we also observed that the *lytS* knockout mutant was cleared more readily from the vaginal tract compared to its wild-type counterpart. Interestingly, lower levels of proinflammatory cytokines were found in the serum of mice infected with the *lytS* mutant. Our results demonstrate that the LytSR TCS plays a key role in GBS tissue invasion and pathogenesis, and persistence of mucosal colonisation.

**Importance:** *Streptococcus agalactiae (Group B strep, or GBS)* is a common commensal of the female urogenital tract and one of WHO’s priority pathogens. The bacterium has evolved mechanisms to adapt and survive in its host, many of which are regulated via two-component signal transduction systems (TCSs), however, the exact contributions of TCSs towards GBS pathogenicity remain largely obscure.

We have constructed a TCS *lytS-*deficient mutant in a CC-17 hypervirulent GBS clinical isolate. Using murine models, we showed that LytSR regulatory system is essential for vaginal colonisation via promoting biofilm production. We also observed that *lytS* deficiency led to significantly attenuated virulence properties and lower levels of proinflammatory cytokines in blood. Our findings are of significant importance in that they unveil a previously unreported role for LytSR in GBS and pave the way towards a better understanding of its ability to transition from an innocuous commensal to a deadly pathogen.

## Introduction

*Streptococcus agalactiae*, commonly known as Group B *Streptococcus* (GBS), is an opportunistic pathogen that commonly colonises the gastrointestinal and genitourinary tracts of healthy adults, the elderly and pregnant women [1–3]. Under given circumstances, GBS can cause life-threatening invasive diseases such as pneumonia, sepsis and meningitis especially in vulnerable groups such as neonates [1]. To date, ten GBS serotypes have been described based on the type of capsular polysaccharide, with serotypes Ia, Ib, II, III and V being the most common causes of life-threatening and self-limiting infections [4]. Multilocus sequence typing (MLST) system was developed to further classify GBS into clonal complexes (CCs) [5]. Certain CCs were found to be commonly associated with invasive disease e.g., CC-17, while others are mainly presented as colonising strains e.g., CC-1 and CC-19 [5, 6]. In a recent report, the genomic sequence analysis of nearly 2,000 GBS carriage and invasive disease-associated isolates originating from different hosts (human and animal) and five different countries led to the identification of a hundred accessory and core genes associated with the hypervirulent CC-17. Amongst them, the proton-dependent oligopeptide transporter (POT) family and iron complex transport system permease protein FeuC were deemed particularly important [7].

GBS possesses a number of virulence factors including adhesins such as Srr1/Srr2, the fibrinogen-binding proteins (FbsA, FbsB and FbsC), the laminin binding protein (Lmb), the surface-associated serine protease ScpB or C5a peptidase, the plasminogen binding surface protein (PbsP), the pili (proteins PilA, PilB and PilC), the GBS immunogenic bacterial adhesin (BibA) and the hyaluronidase HylB (reviewed in [8, 9]). The polysaccharide capsule encoded by the *cps* operon participates to GBS resistance to complement deposition, opsonisation and phagocytosis [10, 11], while the β-hemolysin/cytolysin (β-H/C), also referred as the “hemolytic pigment” or Granadaene, is responsible for the hemolytic activity of GBS and is encoded by the *cyl* operon [12, 13]. In CC-17 in particular, the adhesins Srr2 and HvgA, and the pilus protein PI-2b are known to play an important roles in host cell adhesion, colonisation, and tolerance to phagocytosis [14–16]. While all these GBS virulence factors are well documented, their genetic regulation remains largely unexplored.

Of the regulatory factors known to be essential [6], two-component regulatory systems (TCSs) are major contributors to bacterial responses to environmental changes through regulation of gene expression in response to stimuli [17, 18]. TCSs consist of a sensor histidine kinase that once activated by an external signal or stress conditions, autophosphorylates its cytoplasmic domain and relays the phospho-group to the transcriptional response regulator that in turn regulates the expression of target genes [18]. The size and complexity of the regulons for TCS are highly variable, ranging from a few genes to hundreds of genes [19]. To date, around 20 TCSs have been described in GBS, some of which have been shown to play important roles in bacterial pathogenesis [20]. Examples include the CovRS, in which deletion mutants have shown different effects on virulence depending on the infection model used [20–22] and HssRS, a regulator of heme transport which has been shown to be critical during systemic infection in a mouse model [23]. Furthermore, a number of TCSs in GBS are known to be important for regulation of colonisation, adhesion and resistance to antimicrobials [20]. Genome analysis of *S. agalactiae* has also identified orthologs of TCSs known to participate to the virulence in other species, however, their role in *S. agalactiae* is yet unknown. One such example is LytSR [24].

The LytSR two-component system is composed of the sensor LytS and the regulator LytR. LytSR is widely conserved across species and has been shown to be important for virulence-associated phenotypes such as biofilm formation and resistance against host cationic antimicrobial peptides (CAMPs) [25]. In different Staphylococcal species, lack of LytSR has been shown to alter biofilm formation capabilities potentially by altered autolysis [24, 26, 27], while deletion of *lytS* led to a significantly increased susceptibility to membrane-damaging CAMPs in both *in vitro* and *in vivo S. aureus* infection models [25]. The direct regulatory function of LytSR has been linked to cell autolysis and metabolism in different species, as the LytSR system has been shown to sense both changes in membrane potential and extracellular metabolic signals such as pyruvate, glucose and oxygen [28]. From studies in *S. aureus*, *Staphylococcus epidermidis*, *Streptococcus mutans* and *Bacillus subtilis,* it is established that LytSR controls the expression of the downstream *lrgAB* operon which encodes a pore-forming holin [29]. The actual role of LrgAB in not fully established; it may be important for the regulation of murein hydrolase activity similar to other well-characterised holins, but recent research has also indicated that LrgAB is transporting metabolic by-products such as pyruvate [29, 30]. Regulation of LrgAB by LytSR could thus also be used by the bacteria to integrate extracellular signals with the metabolic state of the cell [28]. Furthermore, transcriptional analyses have indicated that LytSR have a global regulatory role. Microarray analysis in *S. aureus* revealed that removal of *lytSR* affected the expression of 267 genes encoding proteins involved in a variety of functions such as carbohydrate, energy and nucleotide metabolism [27]. Similarly, transcriptional analysis of *lytSR* mutants in *S. mutans* and *S. epidermidis* showed equivalent effects on gene expression [31, 32]. However, the role of LytSR two-component system in GBS pathogenesis remains largely unknown. Here, we aimed to investigate the impact of *lytS* deletion *in vitro* on GBS autolysis, hemolysis, cell adhesion and biofilm formation, as well as on *in vivo* on mucosal colonisation and invasive properties.

## Results

### Phenotypic characterisation of *lytS* deletion in GBS

To investigate the contribution of LytSR in GBS virulence, we deleted the *lytS* gene (encoding the sensor of the system) in a serotype III CC-17 clinical isolate (strain HQ199) to generate the isogenic *lytS* deficient strain HQ199Δ*lytS*. We first compared the growth of the Δ*lytS* mutant to its wild-type (WT) counterpart in planktonic cultures *in vitro*. Both strains showed similar growth over 24 hours in Todd-Hewitt broth (Fig. 1A). We also monitored the survival of both strains in human whole blood, this experimental setup serving as an *ex vivo* assay to mimic the *in vivo* blood environment. Both WT and Δ*lytS* isolates showed similar survival rates in human whole blood over a 48-hour period (Fig. 1B). These results indicate that the absence of the *lytS* gene did not affect bacterial growth or survival *in vitro*.

**Figure 1.**
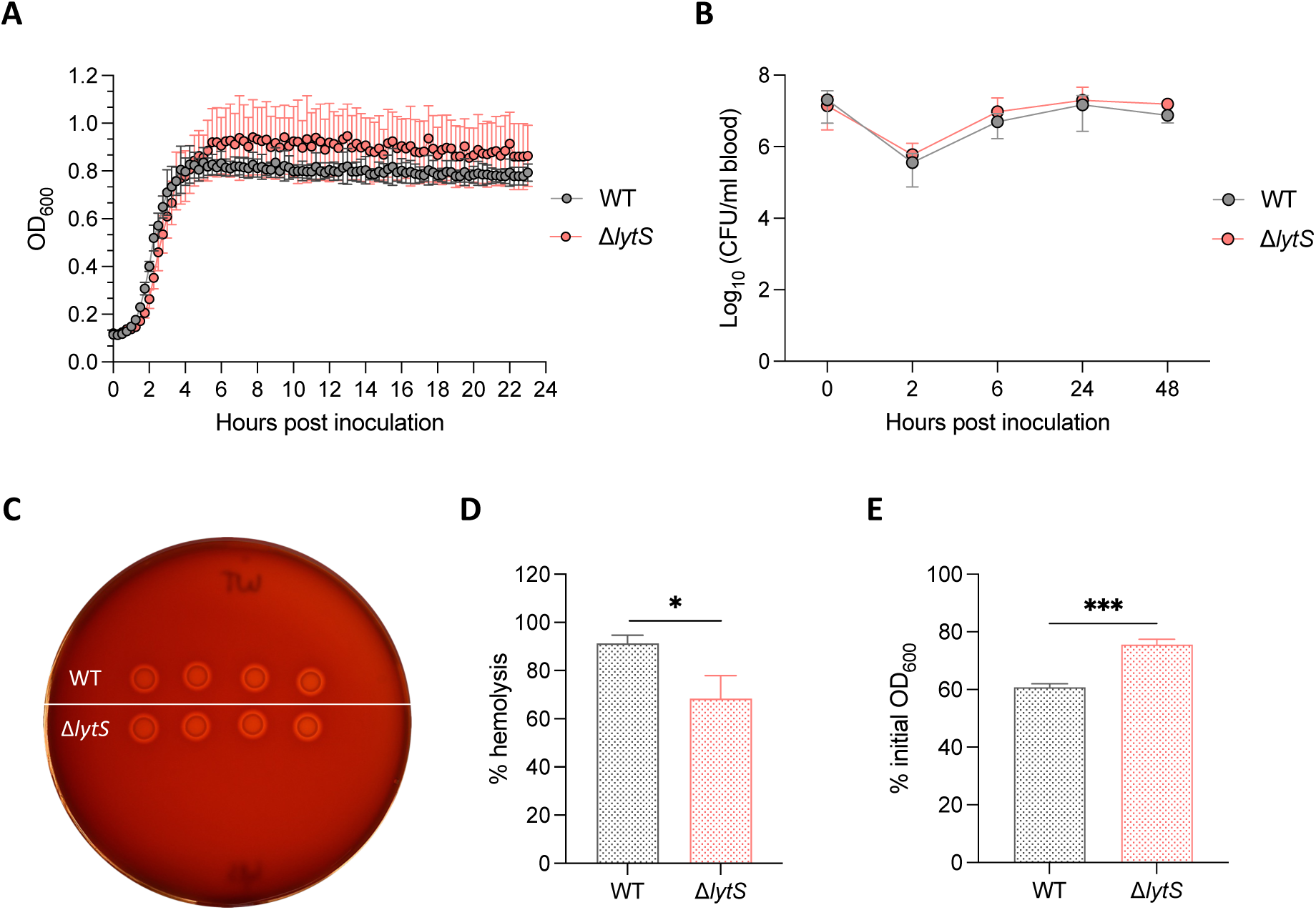
Phenotypic characterisation of GBS wild-type and the isogenic Δ*lytS* strains. (**A**) Bacterial growth of GBS strain HQ199 wild-type (WT) and the isogenic Δ*lytS* mutant was determined in Todd-Hewitt broth at 37°C and 5% CO_2_. Optical density at 600 nm (OD_600_) was measured for 24 hours. Data represent the mean ± SD of three independent experiments. (**B**) Bacterial survival in blood. Bacterial strains were grown in human whole blood at 37°C for 48 hours and viable bacteria were enumerated. Data represent the mean ± SD of three independent experiments. (**C**) Observation of β-hemolysis on sheep blood agar base medium. Bacteria were grown at 37°C overnight. (**D**) Hemolytic activity. Bacterial strains were incubated in fresh sheep red blood cells. After incubation for 30 min, samples were centrifuged, and the levels of hemoglobin released in the supernatant was determined by measuring the optical density at OD_420_. Data represent the mean ± SD of three independent experiments. (**E**) Rates of bacterial autolysis. Bacterial strains were grown at an OD_600_ = 1 prior incubation in 0.01% Triton-X-100 at 37°C with shaking. After incubation for 120 mins, OD_600_ was measured and converted as the percent of the initial OD_600_ reading. Data represent the mean ± SD of three independent experiments. Statistical analysis was performed using the unpaired *t* test. *p < 0.05 and ***p < 0.001.

Next, we analysed the hemolytic activity of both WT and Δ*lytS* strains. Macroscopic observation on blood agar base medium showed similar β-hemolysis for both isolates (Fig. 1C), however quantification of hemolysis showed significantly decreased hemolytic activity in the *lytS* mutant compared to the WT strain (Fig. 1D), suggesting that *lytS* deficiency impaired the ability of GBS to lyse red blood cells. Similarly, we investigated the impact of *lytS* deficiency on GBS autolysis. Triton X-100-induced autolysis assay showed that the Δ*lytS* mutant displayed a significantly lower rate of autolysis compared to the WT strain (Fig. 1E and S1), which suggests that *lytS* deficiency impaired autolysis in GBS.

### *lytS* deficiency alters GBS colonisation in the genital tract

The genital tract is a common tissue niche for GBS colonisation [33]. Hence, we explored whether and how *lytS* deficiency impacted on vaginal colonisation *in vivo*. One day prior to infection with GBS, female Balb/c mice were treated with 17β-estradiol hormone to mimic the estrus cycle [34]. The following day, mice were intravaginally colonised with 1 × 10^6^ CFU of GBS WT or Δ*lytS* strains, and the level of bacterial colonisation was quantified in vaginal, cervical, and uterine tissues over a 21-day period. Both WT and Δ*lytS* strains established stably and at equivalent levels in the vagina and cervix of mice during the first seven days following infection (Fig. 2A and B), and both isolates were also detectable in the uterus at similar levels since day 1 post infection (Fig. 2C), suggesting that both GBS isolates migrated rapidly from the vagina to the uterus. Interestingly, at day 14 post infection, mice infected with the Δ*lytS* mutant displayed significantly lower CFU counts in the vagina (p= 0.03) while several mice showed no bacteria in cervix or uterus tissues. A total clearance of the mutant strain was observed in all tissues at day 21 post infection (Fig. 2). Conversely, colonisation with the WT strain remained stable and persisted at a density of 10^4^ CFU/ml of tissue in the vagina, cervix and uterus up to day 21 post infection (Fig. 2). These results show that the absence of *lytS* resulted in reduced vaginal colonisation and promoted a more rapid clearance of GBS from the genital tract.

**Figure 2.**
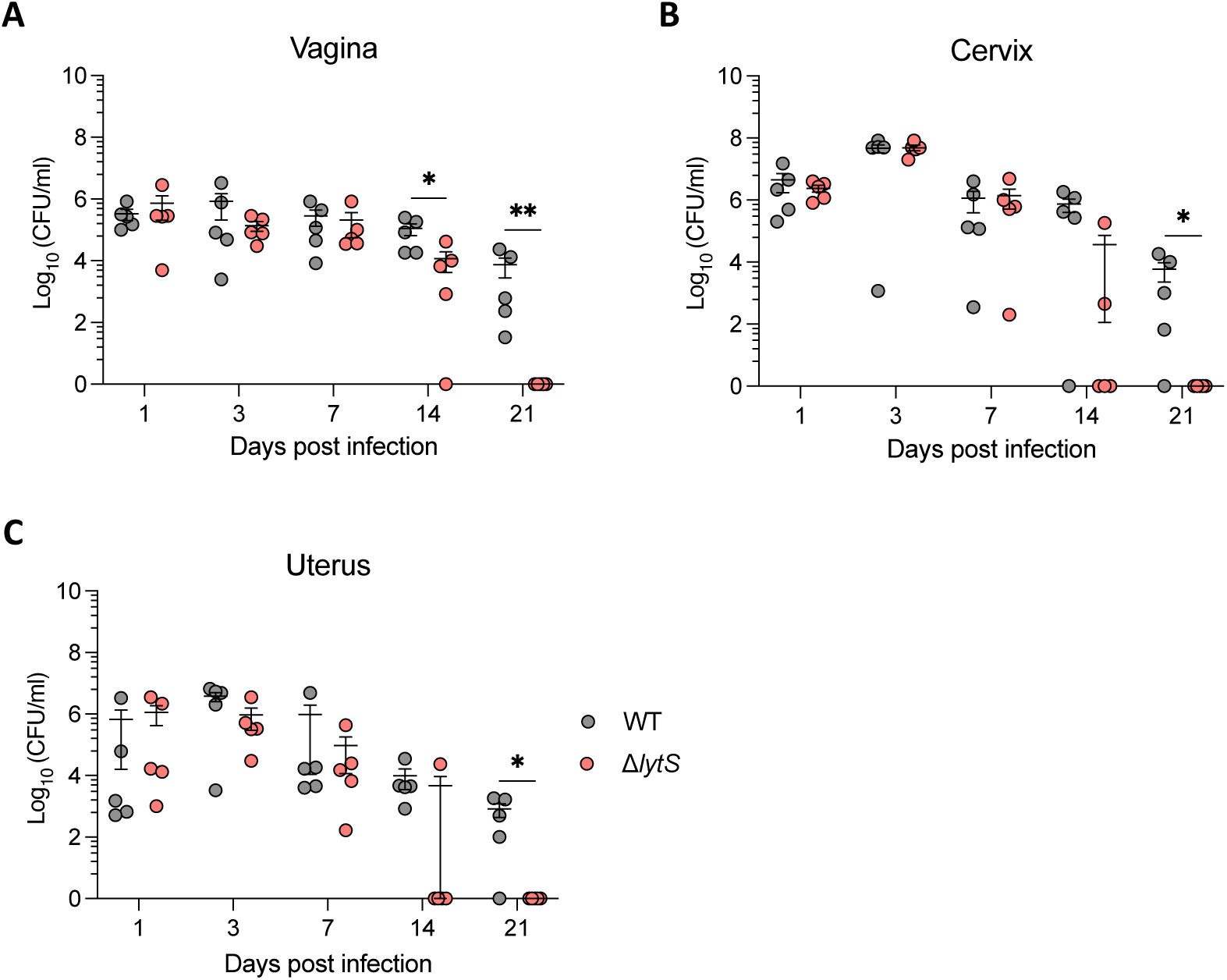
*lytS* deficiency impairs GBS colonisation and persistence in the genital tract of mice. Female Balb/c mice (n = 30 per group, five mice per day) were infected intravaginally with 1 × 10^6^ CFU of GBS HQ199 wild-type (WT) or the isogenic Δ*lytS* strains. Levels of bacterial colonisation in the vagina (**A**), cervix (**B**) and uterus (**C**) were quantified at days 1, 3, 7, 14 and 21 post infection. Each dot represents a single mouse and error bars represent the mean ± SEM in each time point. Statistical analysis between the two groups in each day was performed using the Mann-Whitney U test. *p < 0.05 and **p < 0.01.

We next examined whether the impaired ability of the Δ*lytS* mutant to colonise the genital tract of mice was due to an alteration of its capability to adhere and/or invade the vaginal epithelium and produce biofilm. We performed in vitro adherence and invasion assays using the human vaginal epithelial VK2/E6E7 cell line. The Δ*lytS* mutant showed slightly reduced adhesion and invasion to epithelial cells compared to its WT counterpart (Fig. S2A and B), albeit this was not statistically significant. In addition, the Δ*lytS* mutant produced significantly less biofilm than the WT strain *in vitro* (p < 0.01) (Fig. S2C). Taken together, these results suggest that *lytS* deficiency impaired the ability of GBS to colonise the genital tract of mice by reducing its biofilm production and altering its cell adhesion and invasion properties.

### LytSR plays a key role in GBS virulence and dissemination *in vivo*

We next sought whether the deletion of *lytS* contributes to GBS virulence and pathogenesis *in vivo* using a mouse sepsis model of GBS infection. All female Balb/c mice intravenously infected with the WT strain displayed 100% mortality within 36 hours of infection, while all mice infected with the Δ*lytS* mutant survived the infection (Fig. 3A). The bacterial density in tissue quantified at time of death showed that infection with the Δ*lytS* mutant was characterised by significantly reduced bacterial loads in blood, lung and brain tissues compared to mice infected with the WT parent strain (Fig. 3B). These results suggest that the LytSR TCS played a key role in GBS virulence during sepsis.

**Figure 3.**
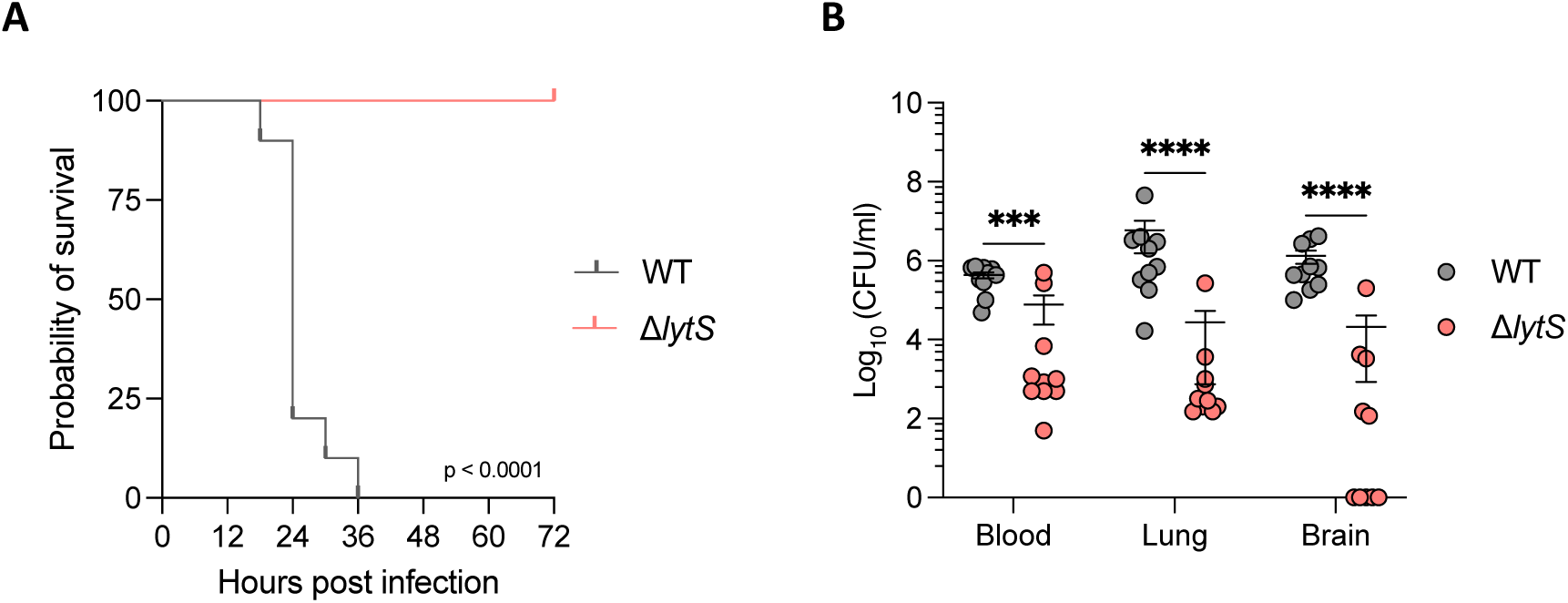
*lytS* deficiency reduces GBS virulence and dissemination in a mouse sepsis model. (**A**) Kaplan-Meier curve representing probability of survival of female Balb/c mice (n = 10 per group) infected intravenously with 1 × 10^8^ CFU of GBS HQ199 wild-type (WT) or the isogenic Δ*lytS* strains. Statistical analysis was performed using the Log-rank (Mantel-Cox) test. (**B**) Levels of bacterial loads in blood, lung and brain tissues following euthanasia of mice. Each dot represents a single mouse and error bars represent the mean ± SEM. Statistical analysis was performed using the Mann-Whitney U test. ***p < 0.001 and ****p < 0.0001.

To further investigate this, we next analysed the kinetics of dissemination of both mutant and WT isolates during sepsis. Following intravenous administration of bacteria, both strains colonised blood at equivalent levels (30min post-challenge Fig 4A). However, between 24 and 32 hours post infection, the bacterial burden in Δ*lytS-*infected mice significantly decreased in blood, and all tissues compared to mice infected with the WT isolate (Fig. 4A-C). In the kidney, a significantly lower bacterial density was observed in Δ*lytS*-infected mice as early as 6 hours post infection and this pattern persisted until 32 hours post infection (Fig. 4D). The largest differences were observed in brain tissue where WT-infected animals showed a rapid and gradual increase in CFU counts, going from 10^2^ CFU/ml of tissue at 30 min post infection to 10^6^ CFU/ml of tissue at 32 hours post infection (Fig. 4E). In contrast, no viable bacteria were detected in the brain of mice infected with the Δ*lytS* mutant until 32 hours post infection, where two mice were colonised at a density inferior to 10^2^ CFU/ml of tissue (Fig. 4E). Altogether, these results indicate that *lytS* deficiency impaired the dissemination of GBS from blood to other tissues, especially to the brain, and resulted in a more rapid bacterial clearance.

**Figure 4.**
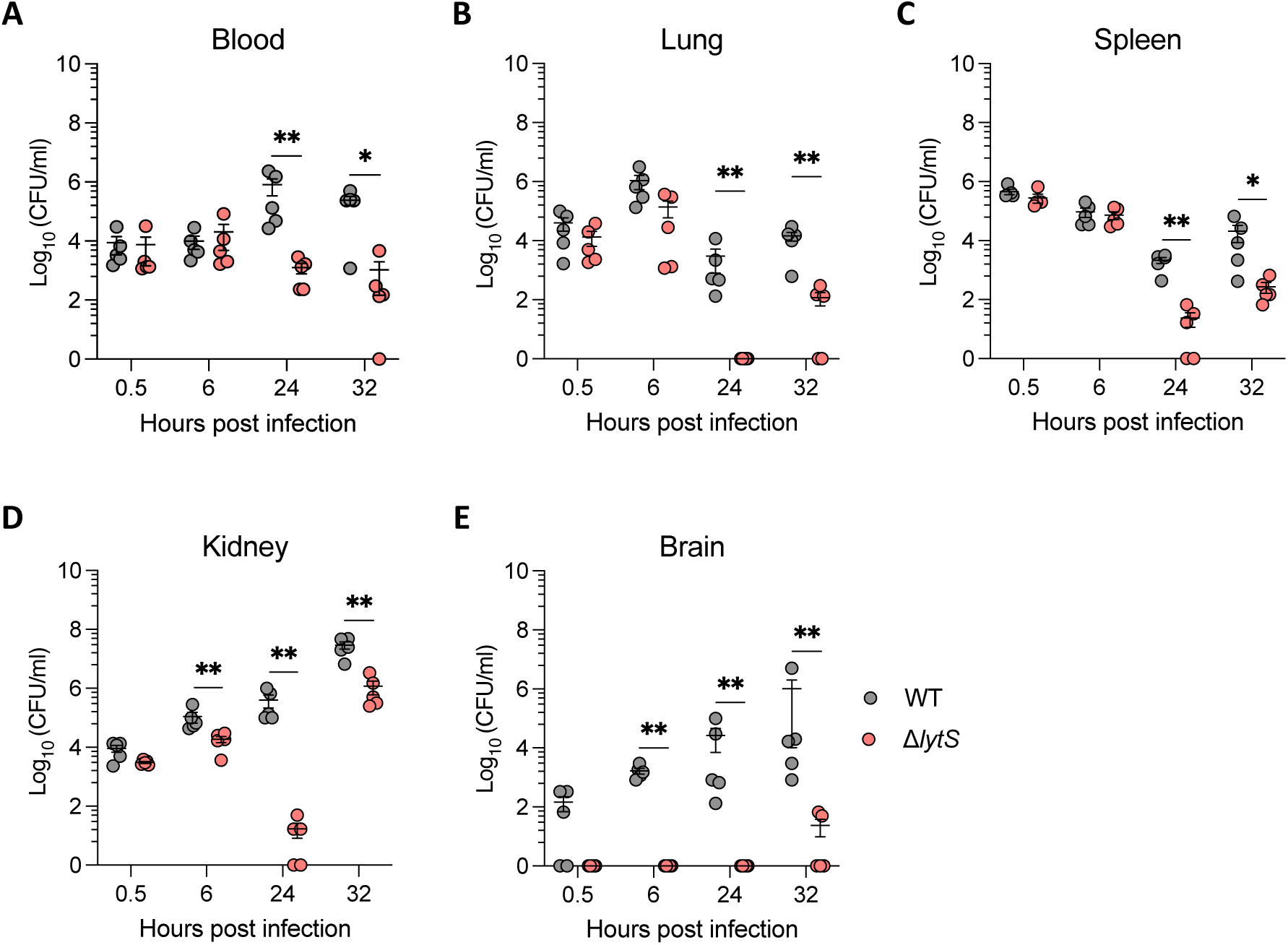
*lytS* deficiency prevents bacterial dissemination and brain invasion in vivo in a mouse sepsis model. Female Balb/c mice (n = 20 per group, five mice per day) were intravenously infected with 1 × 10^8^ CFU of GBS HQ199 wild-type (WT) or the isogenic Δ*lytS* strains. Levels of bacterial loads in blood (**A**), lung (**B**), spleen (**C**), kidney (**D**) and brain (**E**) tissues were quantified at 0.5, 6, 24 and 32 hours post infection. Each dot represents a single mouse and error bars represent the mean ± SEM in each time point. Statistical analysis between the two groups in each time point was performed using the Mann-Whitney U test. *p < 0.05 and **p < 0.01.

### *lytS* deficiency results in attenuated host inflammatory responses

As the absence of LytSR resulted in significant attenuation of GBS virulence, we examined the host immune response during sepsis by quantifying the levels of the proinflammatory cytokines IL-1β, IL-2, IL-6, IL-17A/F, MIP-3α, KC/GRO and TNF-α in the serum of mice infected with the WT or Δ*lytS* strains. Up to 6 hours post infection, no differences were observed between the 2 groups of mice. At 24 and 32 hours post infection, we measured significantly lower levels of IL-1β, IL-6, MIP-3α and KC/GRO in mice infected with the Δ*lytS* mutant, as well as for IL-2 cytokine at 32 hours post infection (Fig. 5A-C and E-F). Transient higher levels of IL-17A/F and TNF-α were observed at 30 min post infection in the Δ*lytS* mutant group, but at 24 and 32 hours post infection, levels of both cytokines were found to be significantly lower compared to the WT-infected animals (Fig. 5D and G). Interestingly, differences in cytokine patterns closely coincided with the kinetics of bacterial loads observed in tissues (Fig. 4). Altogether, these results indicate that attenuation of GBS virulence due to the absence of LytSR also reduced its ability to induce host inflammatory responses during infection.

**Figure 5.**
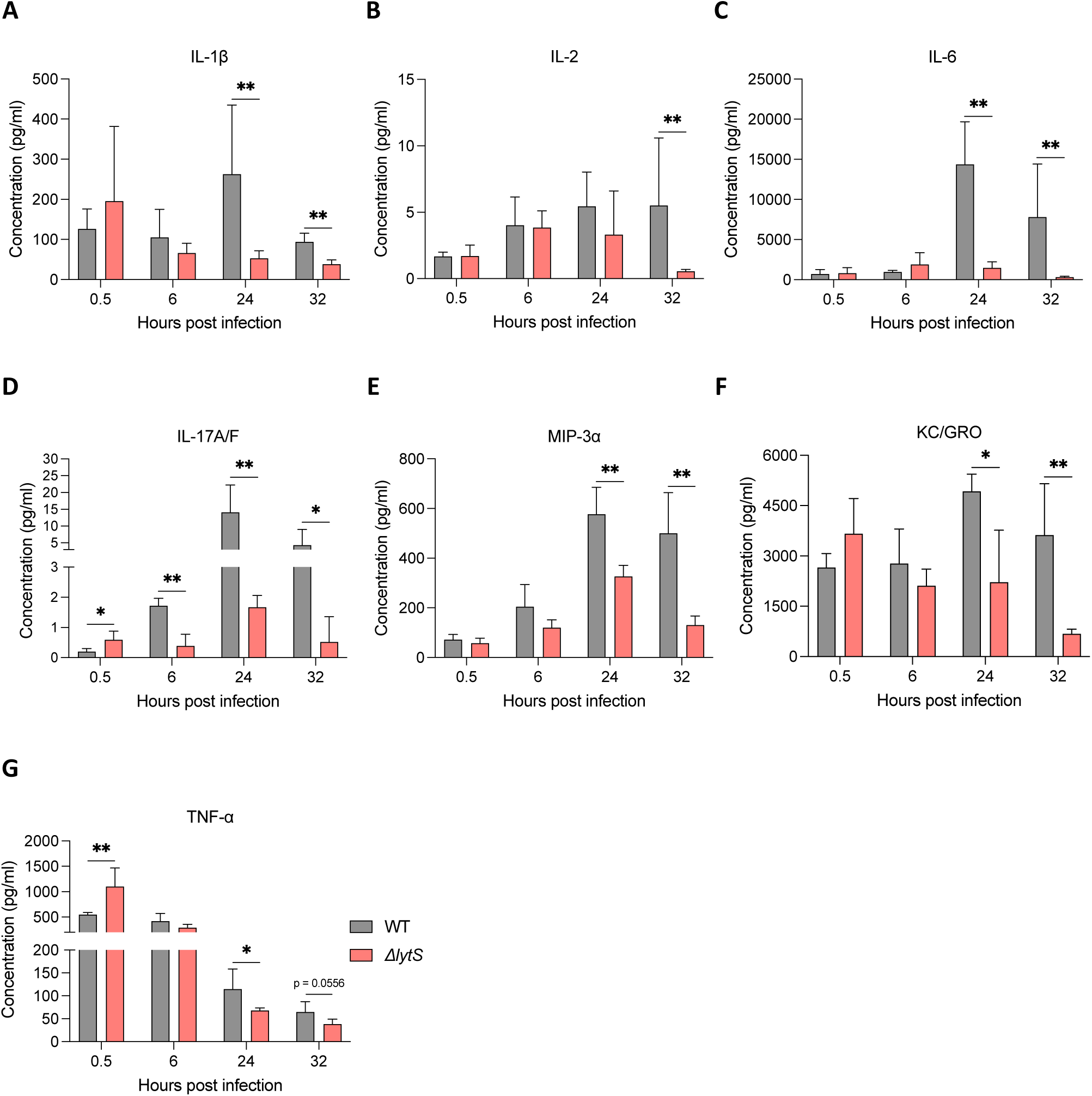
*lytS* deficiency impacts proinflammatory host responses during GBS sepsis. Female Balb/c mice (n = 20 per group, five mice per day) were intravenously infected with 1 × 10^8^ CFU of GBS HQ199 wild-type (WT) or the isogenic Δ*lytS* strains. Levels of proinflammatory cytokines IL-1β (**A**), IL-2 (**B**), IL-6 (**C**), IL-17A/F (**D**), MIP-3*α* (**E**) KC/GRO (**F**) and TNF-α (**G**) were quantified in the serum of mice at 0.5, 6, 24 and 32 hours post infection. Data represent the mean ± SD in each time point. Statistical analysis between the two groups in each time point was performed using the Mann-Whitney U test. *p < 0.05 and **p < 0.01.

## Discussion

*S. agalactiae* is a natural commensal of the genitourinary and gastrointestinal tracts, yet it is known to be an important cause of invasive disease in neonates and adults. Around 20 TCSs are encoded in GBS, which represents almost twice as many as other related Streptococcal species such as *S. pneumoniae* or *S. pyogenes*, both of which possess only 13 TCSs within their entire genome [20, 35–37]. This suggests that GBS has a strong and finely tuned capacity for sensing and responding to multiple environmental stimuli [20]. The LytSR TCS was first identified in GBS in 2002 [17]. However, to the best of our knowledge, its role in GBS virulence and pathogenesis has never been explored experimentally in clinically relevant models. Here we show, using an isogenic *lytS*-deficient mutant (where *lytS* encodes the sensor histidine kinase of the TCS), that LytSR has a major role to play during GBS colonisation and pathogenesis *in vivo*.

Our results show that the LytSR TCS plays a significant role in GBS virulence during sepsis, whereby all mice infected with the WT strain succumbed to infection within 36 hours post infection, while all animals infected with the *lytS* mutant survived infection. Interestingly, the *lytS-*deficient strain showed a decrease in hemolytic activity, a feature that may have contributed to its reduced virulence [38–41]. Infection with the Δ*lytS* mutant was associated with a significant reduction of viable bacteria in blood, lung, spleen, kidney and, most dramatically, in the brain. Thus, LytSR would seem to be involved in *S. agalactiae* survival and proliferation in blood and tissue during different infection stages, due to reduced bacterial survival in the absence of *lytS* in vivo. Notably, the dramatic reduction of bacterial load in the brain (Fig. 4E), suggests that the Δ*lytS* mutant may have a reduced capacity to cross the blood brain barrier (BBB). Proinflammatory cytokines are known to disrupt tight junctions and increase permeability to pathogens at the BBB [42–44]. Barichello et al. showed that levels of CINC-1 (also known as KC/GRO or CXCL1), IL-1β, IL-6 and TNF-α were increased in the hippocampus of neonate rats during GBS meningitis in association with BBB breakdown [45]. Cytokines IL-2, IL-17A and the chemokine MIP-3α (CCL20) have also been shown to be associated with bacterial meningitis and promote BBB disruption [46–49]. Similarly, previous studies have shown that the GBS hemolytic pigment facilitates the penetration of GBS into the BBB and promotes proinflammatory responses in microvascular endothelial cells [21, 38]. Altogether, these previous observations are in line with our results presented here, showing that reduced dissemination of bacteria into mouse brain tissue upon infection with the Δ*lytS* mutant accompanied by lower productions of proinflammatory cytokines (Fig. 5).

Using a mouse model of vaginal colonisation, we also observed that *lytS* deficiency altered the capacity of GBS to stably establish, proliferate and persist in the genital tract. While WT bacteria showed stable and persisting colonisation in the genital tract of mice over 21 days of infection, the Δ*lytS* mutant isolate was cleared from day 14 post infection and no viable bacteria were detectable in the genital tract on day 21 post infection. This effect could possibly be due to the reduced ability of *lytS-*deficient GBS to adhere to epithelial cells, invade the genital tract and produce biofilms (Fig. S2). Previous studies have shown that biofilm formation could enhance the ability of GBS to colonise its host and induce invasive disease [50–53]. Biofilm production is thought to be closely related to bacterial autolysis, on the basis that extracellular DNA released following cell lysis contributes to biofilm formation [54–56]. As such, reduced autolysis observed in the *lytS* mutant strain lacking LytSR may be associated with its reduced capacity to for biofilm production.

To the best of our knowledge, our study is the first to report that the LytSR two-component regulatory system plays a key role in GBS virulence and pathogenesis. Our result further suggests that the importance of LytSR is due to its effect on cell adhesion, biofilm production, colonisation and invasiveness. In other species, the LytSR system senses changes in membrane potential as well as extracellular metabolic signals, such as the presence of oxygen, pyruvate or glucose to regulate expression of a holin and potentially hundreds of other genes [28, 57]. Further studies of gene regulation both in vitro and in vivo are needed to characterise how LytSR TCS affects gene expression in *S. agalactiae* during the course of infection, and to understand whether the altered pathogenesis is resulting from direct regulation of known virulence factors by LytSR or whether novel mechanisms are at play. This may pave the way towards the finding of novel and more efficient strategies to prevent GBS colonisation and invasive disease, particularly in susceptible populations.

## Materials and Method

### Bacterial strains and growth conditions

*S. agalactiae* strain HQ199 is a serotype III CC-17 clinical isolate isolated from the vaginal tract of a pregnant woman during an antenatal screening at the Liverpool Women’s Hospital. The isogenic MB-HQ199Δ*lytS* mutant was constructed as described further. GBS was routinely grown on blood agar base (BAB) medium or in Todd-Hewitt broth (THB) at 37°C. For growth assay in nutritive medium, a 2 ml culture volume containing 200 CFU of GBS was grown in THB supplemented with 20% fetal bovine serum in 24-well U-bottom microplates (Greiner). Bacterial growth was monitored every 15 min for a period of 24 hours by spectrophotometry measuring the optical density at 600 nm (OD_600_) using a FLUOstar Omega microplate reader (BMG Labtech). For blood survival assay, 300 μl of heparinised human blood were inoculated with 50 CFU of GBS and incubated at 37°C at 180 rpm. At specified time points, 100 μl of the mixture were serially diluted and dilutions were plated onto Todd-Hewitt agar medium and incubated at 37°C overnight. Viable bacteria were enumerated and expressed as CFU/ml blood. Photographs of bacterial strains grown on sheep BAB medium were captured with a Nikon D500 camera with a 16-80 mm lens.

### Construction of Δ*lytS*-deficient mutant

The isogenic *lytS* knockout mutant in *S. agalactiae* strain HQ199 was constructed by replacing *lytS* with the *aad9* cassette (conferring resistance to spectinomycin) through allelic replacement. Briefly, the *aad9* cassette and the flanking regions of *lytS* (∼1000 bp each) were amplified by PCR. The PCR products were stitched together in a second round of PCR generating the Δ*lytS::aad9* deletion fragment which was then ligated into pHY304 vector, generating pHY304-Δ*lytS::aad9* plasmid in which the deletion fragment provided a mean of selection of transformants. pHY304-Δ*lytS::aad9* was transformed into competent GBS cells. Transformants were recovered on Todd-Hewitt agar medium supplemented with spectinomycin (Sigma; 200 μg/ml). The authenticity of *lytS* deletion in the selected transformant MB-HQ199Δ*lytS* was verified by PCR and Sanger sequencing.

### Hemolytic activity assay

Bacteria grown in THB at 37°C to an OD_600_ = 0.4 were centrifuged at 3,000 × g for 10 min, washed with 1X Phosphate Buffered Saline (PBS) and resuspended in 1 ml PBS. In a 96-well U-bottom microplates, 100 μl per well (10^8^ CFU) of the bacterial suspension were placed in the first well, and serial twofold dilutions in PBS were performed across the plate, each in a final volume = 100 μl. An equal volume of 4% fresh sheep red blood cells (RBCs) (TCS Biosciences) washed once and resuspended in PBS, was added to each well. The plates were sealed and incubated at 37°C for 1 hour. PBS alone and 0.1% sodium dodecyl sulfate (SDS) were used as negative and positive controls for hemolysis, respectively. After incubation, the plates were centrifuged at 3000 × g for 10 min to pellet unlysed RBCs, and 100 μl of the supernatant were transferred into a 96-well flat-bottom microplates. Hemoglobin release was determined measuring the optical density at 420 nm (OD_420_), and percent hemolysis for the GBS-treated wells was determined relative to SDS-treated positive and PBS-treated negative controls. Three experiments were performed in triplicate.

### Autolysis assay

Triton X-100 induced autolysis assay was performed as described previously with the following modifications [58, 59]. Briefly, bacteria grown overnight in THB at 37°C to an OD_600_ = 1 were centrifuged at 3,600 × g for 10 min. Bacterial pellets were resuspended in PBS and adjusted to an OD_600_ = 1 in 1 ml PBS containing 0.1% Triton X-100 (Sigma). The initial OD_600_ was measured (T_0_). Bacterial suspensions were then incubated at 37°C with shaking (180 rpm), and the OD_600_ was recorded every 15 min for a period of 2 hours. Autolysis was determined as the percent of the initial OD_600_ reading at T_0_. Three experiments were performed in triplicate.

### Biofilm assay

Biofilm production was determined by crystal violet (CV) staining for measuring attached biofilm as described previously by Patras et al [58]. Bacteria grown in THB at 37°C overnight were centrifuged and resuspended in PBS. Cultures were adjusted at an OD_600_ = 0.1 and 1 ml of each culture was distributed into 24-well flat-bottom plates that were incubated at 37°C for 24 hours. After incubation, wells were washed twice with 200 μl PBS to remove planktonic non-adherent cells. Wells were dried for 1 hour at room temperature prior staining with 200 μl of 0.25% CV for 45 min at 55°C. Wells were then washed three times with dH_2_O, and 200 μl of 95% ethanol were added to wells before incubation for 30 min at room temperature. The OD_600_ was measured and wells containing 95% ethanol were used as blank controls. Three experiments were performed in triplicate.

### Adhesion and invasion assays

Bacterial adhesion and invasion were tested by culturing human vaginal epithelial VK2/E6E7 cells in Dulbecco modified Eagle medium (DMEM) supplemented with 10% FBS and 1% penicillin (10,000 U/ml) and 1% streptomycin (10 mg/ml), to > 90% confluency in 24-well plates at 37°C in 5% CO_2_. Then, 1 × 10^7^ bacteria in 1 ml DMEM were incubated at 37°C in 5% CO_2_ for 60 min for adhesion or 2 hours for invasion. After incubation, wells were washed 3 times with PBS. Adherence was tested by trypsinising cells with 100 μl 0.025% trypsin-0.53 mM EDTA, adding 900 μl PBS, and plating samples serially diluted in PBS on BAB medium for determination of the numbers of CFU/well. Invasion assay mixtures were further incubated for 2 hours with 5 µg penicillin and 100 µg gentamicin at 37°C and 5% CO_2_. Wells were washed three times, and cells were trypsinised with 200 μl 0.025% trypsin-0.53 mM EDTA. Cells were lysed with 400 μl 0.1% Triton X-100, and samples were plated on BAB medium to determine the numbers of CFU/well. Three experiments were performed in triplicate.

### Mouse models of GBS vaginal colonisation and sepsis

This study was conducted in strict accordance with the guidelines outlined by the UK Home Office under the Project License Number PB6DE83DA. All animal experimentations were performed at the Biomedical Services Unit, University of Liverpool and approved by the University of Liverpool Animal Welfare and Ethical Review Body.

In vivo experiments were performed using female BALB/c mice (5-7-week-old) obtained from Charles River Laboratories (Kent, UK). Upon delivery, mice were allowed to acclimatise for seven days prior to use.

For vaginal colonisation, mice were synchronised into the estrus cycle by intraperitoneal administration of 17β-estradiol hormone one day prior to intravaginal infection with 1 × 10^6^ CFU of GBS in 10 μl. At days 1, 3, 7, 14 and 21 post infection, mice were euthanised and vaginal, cervix, and uterus tissues were collected and placed into a sterile Bijou container containing 2 ml PBS and mechanically homogenised for ∼ 5 sec using an IKA Disperser (T 10 basic ULTRA-TURRAX^®^, IKA, Germany). Samples were serially diluted in PBS, plated on BAB medium before incubation at 37°Covernight. CFU/ml of tissue were then enumerated.

For sepsis infection, mice were intravenously infected with 1 × 10^8^ CFU of GBS and they were monitored every 2-3 hours for physical signs of disease using a standard scoring system [60]. Mice were humanely euthanised at predefined time points, i.e., 0.5, 6, 24 and 32 hours post infection or when they were scored ’2+ lethargic’, i.e., minimal movement in the absence of application of finger pressure (for survival assays). Following euthanasia, blood, lung, spleen, kidney and brain tissues were collected. All tissues, except blood, were placed into a sterile Bijou container containing 2 ml PBS and processed as described above to enumerate the number of CFU/ml of tissue. Blood samples collected by tail bleeding at the predefined time points or cardiac puncture following euthanasia in survival assays, were serially diluted in PBS and plated on BAB medium. CFU/ml were then enumerated.

### Cytokines analysis

Blood samples collected by tail bleeding were placed into a sterile tube and immediately centrifuged at 12,000 × g for 5 min. Serum of mice was then carefully collected and placed in a sterile tube before storage at -20°C until use. Quantification of cytokines/chemokines IL-1β, IL-2, IL-6, IL-17A/F, MIP-3α, KC/GRO and TNF-α was performed using a U-Plex cytokine assay (Meso Scale Discovery, Rockville, Md) according to the manufacturer’s instructions.

### Data availability

The genome sequence of *S. agalactiae* strain HQ199 used in this study was deposited in the European Nucleotide Archive (ENA). Accession number: XXX.

### Statistical analysis

Statistical analysis was performed using GraphPad Prism 10® (GraphPad Software, Inc., San Diego, CA). Differences were determined using the unpaired *t* test, the Mann-Whitney U test and the Log-rank (Mantel-Cox) test.

## Author contributions

H.A and M.OB. designed the study. H.A., M.B. and M.OB. performed the experiments. M.OB., N.F and A.K supervised the study. H.A, M.B., M.OB wrote first draft, and along with A.K. edited and corrected the final manuscript. M.K., T.P. and A.C. contributed to the interpretation of data. All authors read and approved the final version of the manuscript.

## Acknowledgements

We would like to acknowledge Prof Kelly Doran, University of Colorado Anschutz Medical Campus, for providing the pHY304 plasmid, as well as Mr Timothy Neal (Microbiology Consultant) and Paul Roberts (Senior Biomedical Scientist) at the Royal Liverpool and Broadgreen University Hospitals NHS Trust, for facilitating access to the *S. agalactiae* HQ199 clinical isolate.

## Funding

We acknowledge funding support from the Prince Sattam bin Abdulaziz University through the project number (PSAU/2023/03/26364) awarded to H.A., the Meningitis Now funding awarded to A.K. and M.OB and the UK Medical Research Council Programme Grant (MR/P011284/1) awarded to A.K.

## Conflict of Interest

The authors declare that the research was conducted in the absence of any commercial or financial relationships that could be construed as a potential conflict of interest.

---

**Supplementary fig S1.**
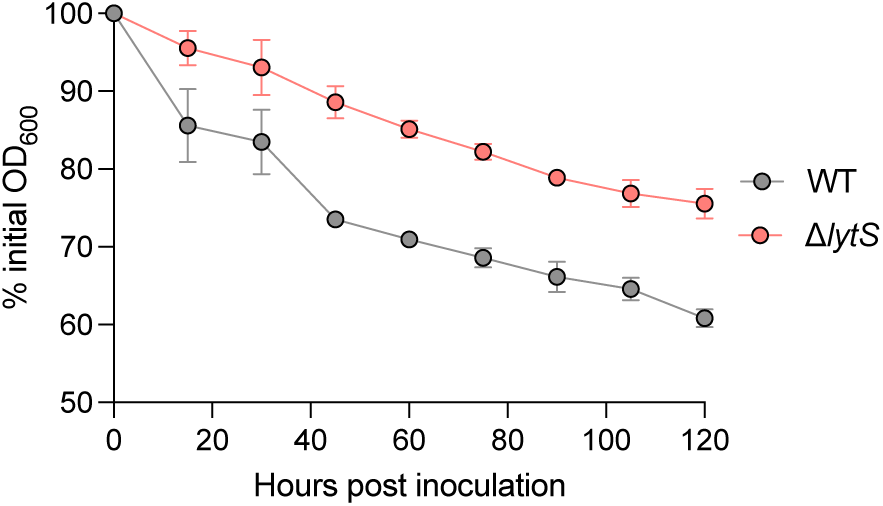
*lytS* deficiency prevents autolysis in GBS. Rates of bacterial autolysis. Bacterial strains were grown at an OD_600_ = 1 prior incubation in 0.01% Triton-X-100 at 37°C with shaking. After incubation, OD_600_ was measured and converted as the percent of the initial OD_600_ reading. Data represent the mean ± SD of three independent experiments.

**Supplementary fig S2.**
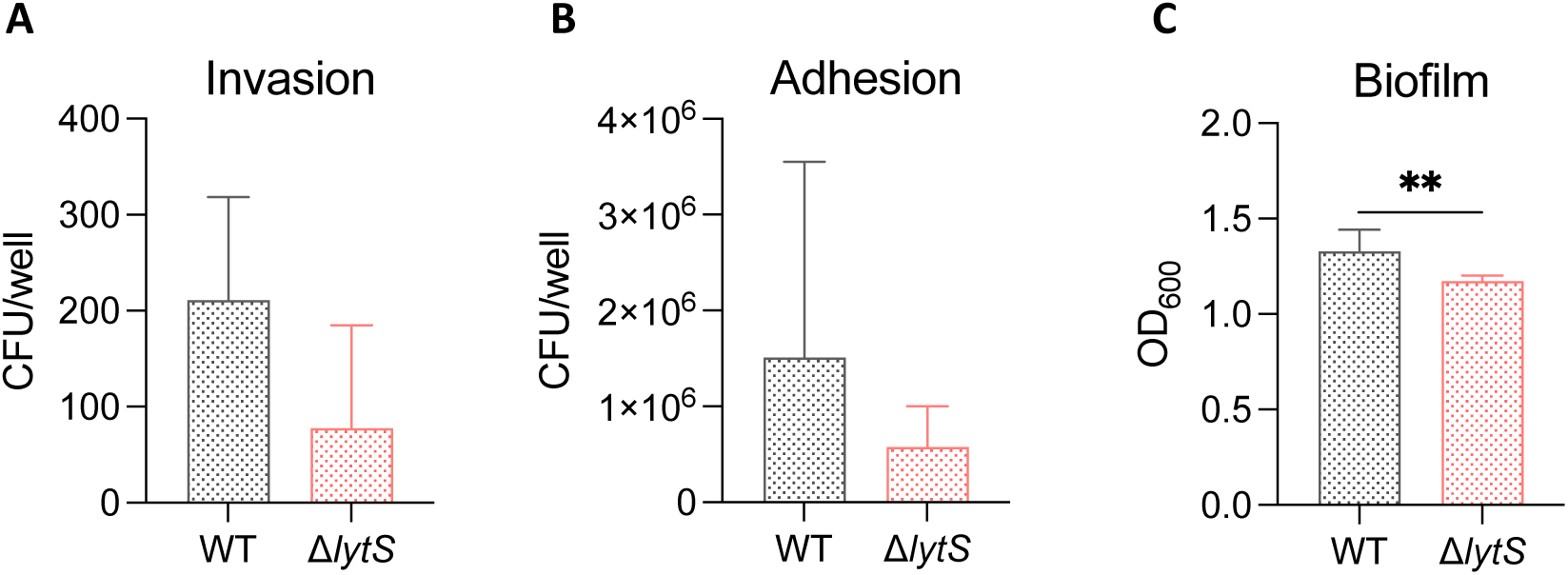
*lytS* deficiency decreases GBS adhesion and invasion on epithelial cells and biofilm formation in vitro. Levels of in vitro bacterial adhesion (**A**) and invasion (**B**) to human vaginal epithelial VK2/E6E7 cells of GBS HQ199 wild-type (WT) and the isogenic Δ*lytS* strains. Adhesion and invasion assays were performed using 1 × 10^7^ CFU of GBS per well. Data represent the mean ± SD of three independent experiments, each performed in triplicate. (**C**) Quantification of the biofilm formation by crystal violet staining. Bacterial strains were grown for 18 hours in Todd-Hewitt broth in 24-well polystyrene plates at 37°C and 5% CO2 and then biofilms were stained with crystal violet (CV). Quantification of the CV staining was performed by spectrophotometry at 600 nm. Data represent the mean ± SD of five independent experiments, each performed in triplicate. Statistical analysis was performed using the Mann-Whitney U test. **p < 0.01.

## Notes

### Competing Interest Statement

The authors have declared no competing interest.

## References

1. Schrag, S.J., et al., Group B streptococcal disease in the era of intrapartum antibiotic prophylaxis. New England Journal of Medicine, 2000. 342(1): p. 15–20.

2. Skoff, T.H., et al., Increasing burden of invasive group B streptococcal disease in nonpregnant adults, 1990–2007. Clinical Infectious Diseases, 2009. 49(1): p. 85–92.

3. Navarro-Torné, A., et al., Burden of invasive group B Streptococcus disease in non-pregnant adults: A systematic review and meta-analysis. Plos one, 2021. 16(9): p. e0258030.

4. Russell, N.J., et al., Maternal colonization with group B Streptococcus and serotype distribution worldwide: systematic review and meta-analyses. Clinical infectious diseases, 2017. 65(suppl_2): p. S100–S111.

5. Jones, N., et al., Multilocus sequence typing system for group B streptococcus. Journal of clinical microbiology, 2003. 41(6): p. 2530–2536.

6. Shabayek, S. and B. Spellerberg, Group B streptococcal colonization, molecular characteristics, and epidemiology. Frontiers in microbiology, 2018. 9: p. 437.

7. Gori, A., et al., Pan-GWAS of Streptococcus agalactiae highlights lineage-specific genes associated with virulence and niche adaptation. Mbio, 2020. 11(3).

8. Liu, Y. and J. Liu, Group B Streptococcus: Virulence Factors and Pathogenic Mechanism. Microorganisms, 2022. 10(12): p. 2483.

9. Rajagopal, L., Understanding the regulation of Group B Streptococcal virulence factors. 2009.

10. Cieslewicz, M.J., et al., Structural and genetic diversity of group B streptococcus capsular polysaccharides. Infection and immunity, 2005. 73(5): p. 3096–3103.

11. Burnham, C.-A.D. and G.J. Tyrrell, Virulence factors of group B streptococci. Reviews and Research in Medical Microbiology, 2003. 14(4): p. 109–118.

12. Whidbey, C., et al., A hemolytic pigment of Group B Streptococcus allows bacterial penetration of human placenta. Journal of Experimental Medicine, 2013. 210(6): p. 1265–1281.

13. Rosa-Fraile, M., S. Dramsi, and B. Spellerberg, Group B streptococcal haemolysin and pigment, a tale of twins. FEMS microbiology reviews, 2014. 38(5): p. 932–946.

14. Lu, B., et al., Microbiological and clinical characteristics of Group B Streptococcus isolates causing materno-neonatal infections: high prevalence of CC17/PI-1 and PI-2b sublineage in neonatal infections. Journal of medical microbiology, 2018. 67(11): p. 1551–1559.

15. Burcham, L.R., et al., Determinants of Group B streptococcal virulence potential amongst vaginal clinical isolates from pregnant women. PloS one, 2019. 14(12).

16. Tazi, A., et al., The surface protein HvgA mediates group B streptococcus hypervirulence and meningeal tropism in neonates. Journal of experimental medicine, 2010. 207(11): p. 2313–2322.

17. Glaser, P., et al., Genome sequence of Streptococcus agalactiae, a pathogen causing invasive neonatal disease. Molecular microbiology, 2002. 45(6): p. 1499–1513.

18. Mascher, T., J.D. Helmann, and G. Unden, Stimulus perception in bacterial signal-transducing histidine kinases. Microbiology and molecular biology reviews, 2006. 70(4): p. 910–938.

19. Rapun-Araiz, B., et al., Systematic reconstruction of the complete two-component sensorial network in Staphylococcus aureus. Msystems, 2020. 5(4): p. 10.1128/msystems.00511-20.

20. Thomas, L. and L. Cook, Two-component signal transduction systems in the human pathogen Streptococcus agalactiae. Infection and immunity, 2020. 88(7): p. e00931–19.

21. Lembo, A., et al., Regulation of CovR expression in Group B Streptococcus impacts blood– brain barrier penetration. Molecular microbiology, 2010. 77(2): p. 431–443.

22. Lamy, M.C., et al., CovS/CovR of group B streptococcus: a two-component global regulatory system involved in virulence. Molecular microbiology, 2004. 54(5): p. 1250–1268.

23. Joubert, L., et al., Visualization of the role of host heme on the virulence of the heme auxotroph Streptococcus agalactiae. Scientific Reports, 2017. 7(1): p. 40435.

24. Brunskill, E.W. and K.W. Bayles, Identification and molecular characterization of a putative regulatory locus that affects autolysis in Staphylococcus aureus. Journal of bacteriology, 1996. 178(3): p. 611–618.

25. Yang, S.-J., et al., Role of the LytSR two-component regulatory system in adaptation to cationic antimicrobial peptides in Staphylococcus aureus. Antimicrobial agents and chemotherapy, 2013. 57(8): p. 3875–3882.

26. Dahyot, S., et al., Role of the LytSR two-component regulatory system in Staphylococcus lugdunensis biofilm formation and pathogenesis. Frontiers in microbiology, 2020. 11: p. 39.

27. Sharma-Kuinkel, B.K., et al., The Staphylococcus aureus LytSR two-component regulatory system affects biofilm formation. Journal of bacteriology, 2009. 191(15): p. 4767–4775.

28. van den Esker, M.H., Á.T. Kovács, and O.P. Kuipers, From cell death to metabolism: Holin-Antiholin homologues with new functions. MBio, 2017. 8(6): p. 10.1128/mbio.01963-17.

29. Ahn, S.-J., et al., Characterization of LrgAB as a stationary phase-specific pyruvate uptake system in Streptococcus mutans. BMC microbiology, 2019. 19(1): p. 1–15.

30. Endres, J.L., et al., The Staphylococcus aureus CidA and LrgA Proteins are functional holins involved in the transport of by-products of carbohydrate metabolism. Mbio, 2022. 13(1): p. e02827–21.

31. Ahn, S.-J., et al., Identification of the Streptococcus mutans LytST two-component regulon reveals its contribution to oxidative stress tolerance. BMC microbiology, 2012. 12(1): p. 1–12.

32. Zhu, T., et al., Impact of the Staphylococcus epidermidis LytSR two-component regulatory system on murein hydrolase activity, pyruvate utilization and global transcriptional profile. BMC microbiology, 2010. 10: p. 1–16.

33. Patras, K.A. and V. Nizet, Group B streptococcal maternal colonization and neonatal disease: molecular mechanisms and preventative approaches. Frontiers in pediatrics, 2018. 6: p. 27.

34. Carey, A.J., et al., Infection and cellular defense dynamics in a novel 17β-estradiol murine model of chronic human group B streptococcus genital tract colonization reveal a role for hemolysin in persistence and neutrophil accumulation. The Journal of Immunology, 2014. 192(4): p. 1718–1731.

35. Chen, S.L., Genomic insights into the distribution and evolution of group B streptococcus. Frontiers in Microbiology, 2019. 10.

36. Gómez-Mejia, A., G. Gámez, and S. Hammerschmidt, Streptococcus pneumoniae two-component regulatory systems: The interplay of the pneumococcus with its environment. International Journal of Medical Microbiology, 2018. 308(6): p. 722–737.

37. Ferretti, J.J., D.L. Stevens, and V.A. Fischetti, Streptococcus pyogenes: basic biology to clinical manifestations [Internet]. 2016.

38. Doran, K.S., G.Y. Liu, and V. Nizet, Group B streptococcal β-hemolysin/cytolysin activates neutrophil signaling pathways in brain endothelium and contributes to development of meningitis. The Journal of clinical investigation, 2003. 112(5): p. 736–744.

39. Ring, A., et al., Group B streptococcal β-hemolysin induces mortality and liver injury in experimental sepsis. The Journal of infectious diseases, 2002. 185(12): p. 1745–1753.

40. Hensler, M.E., et al., Virulence role of group B Streptococcus β-hemolysin/cytolysin in a neonatal rabbit model of early-onset pulmonary infection. The Journal of infectious diseases, 2005. 191(8): p. 1287–1291.

41. Puliti, M., et al., Severity of group B streptococcal arthritis is correlated with β-hemolysin expression. The Journal of infectious diseases, 2000. 182(3): p. 824–832.

42. Yang, R., et al., Blood–Brain Barrier Integrity Damage in Bacterial Meningitis: The Underlying Link, Mechanisms, and Therapeutic Targets. International Journal of Molecular Sciences, 2023. 24(3): p. 2852.

43. De Vries, H.E., et al., The influence of cytokines on the integrity of the blood-brain barrier in vitro. Journal of neuroimmunology, 1996. 64(1): p. 37–43.

44. Zhao, Y., et al., Factors influencing the blood-brain barrier permeability. Brain Research, 2022. 1788: p. 147937.

45. Barichello, T., et al., Oxidative stress, cytokine/chemokine and disruption of blood–brain barrier in neonate rats after meningitis by Streptococcus agalactiae. Neurochemical research, 2011. 36: p. 1922–1930.

46. Kastenbauer, S., et al., Patterns of protein expression in infectious meningitis: a cerebrospinal fluid protein array analysis. Journal of neuroimmunology, 2005. 164(1-2): p. 134–139.

47. Wylezinski, L.S. and J. Hawiger, Interleukin 2 activates brain microvascular endothelial cells resulting in destabilization of adherens junctions. Journal of Biological Chemistry, 2016. 291(44): p. 22913–22923.

48. Coutinho, L.G., et al., Cerebrospinal-fluid cytokine and chemokine profile in patients with pneumococcal and meningococcal meningitis. BMC infectious diseases, 2013. 13: p. 1–9.

49. Xu, B., et al., Meningitic Escherichia coli-Induced Interleukin-17A facilitates blood–brain barrier disruption via inhibiting Proteinase 3/Protease-Activated receptor 2 Axis. Frontiers in Cellular Neuroscience, 2022. 16: p. 814867.

50. Rosini, R. and I. Margarit, Biofilm formation by Streptococcus agalactiae: influence of environmental conditions and implicated virulence factors. Frontiers in cellular and infection microbiology, 2015. 5: p. 6.

51. Noble, K., et al., Group B Streptococcus cpsE is required for serotype V capsule production and aids in biofilm formation and ascending infection of the reproductive tract during pregnancy. ACS infectious diseases, 2021. 7(9): p. 2686–2696.

52. Thomas, L.S. and L.C. Cook, A novel conserved protein in Streptococcus agalactiae, BvaP, is important for vaginal colonization and biofilm formation. Msphere, 2022. 7(6): p. e00421–22.

53. Ho, Y.-R., et al., The enhancement of biofilm formation in Group B streptococcal isolates at vaginal pH. Medical microbiology and immunology, 2013. 202: p. 105–115.

54. Rice, K.C., et al., The cidA murein hydrolase regulator contributes to DNA release and biofilm development in Staphylococcus aureus. Proceedings of the National Academy of Sciences, 2007. 104(19): p. 8113–8118.

55. Qin, Z., et al., Role of autolysin-mediated DNA release in biofilm formation of Staphylococcus epidermidis. Microbiology, 2007. 153(7): p. 2083–2092.

56. Jakubovics, N., et al., Life after death: the critical role of extracellular DNA in microbial biofilms. Letters in applied microbiology, 2013. 57(6): p. 467–475.

57. Lehman, M.K., et al., Identification of the amino acids essential for LytSR-mediated signal transduction in S taphylococcus aureus and their roles in biofilm-specific gene expression. Molecular microbiology, 2015. 95(4): p. 723–737.

58. Patras, K.A., et al., Group B Streptococcus biofilm regulatory protein A contributes to bacterial physiology and innate immune resistance. The Journal of infectious diseases, 2018. 218(10): p. 1641–1652.

59. Oliveira, L., et al., Group B streptococcus GAPDH is released upon cell lysis, associates with bacterial surface, and induces apoptosis in murine macrophages. PloS one, 2012. 7(1): p. e29963.

60. Morton, D., Guidelines on the recognition on pain, distress and discomfort in experimental animals. European Journal of Pharmacology, 1990. 183(4): p. 1583.

